# The major histocompatibility complex: a double-edged sword in fungal disease susceptibility

**DOI:** 10.1101/2024.12.13.628321

**Authors:** Minjie Fu, John A Eimes

**Affiliations:** Research Institute of Basic Sciences, Seoul National University, Seoul 00826, Republic of Korea; School of Biological Sciences, Seoul National University, Seoul 08826, Republic of Korea; University College, Sungkyunkwan University, Suwon 16419, Republic of Korea

**Keywords:** chytridiomycosis, *Batrachochytrium dendrobatidis* (Bd), major histocompatibility complex (MHC), MHC functional modeling, disease susceptibility, allelic expression

## Abstract

Chytridiomycosis, caused by *Batrachochytrium dendrobatidis* (Bd), threatens amphibian populations worldwide. MHC II has been implicated in Bd susceptibility, but the role of MHC in Bd susceptibility remains poorly understood. In this study, we clustered MHC II ß1 alleles into functional supertypes and investigated their diversity in amphibian species with differential susceptibility. Despite sharing similar alpha diversity in MHC II supertypes, the hosts exhibited distinct beta diversity. We demonstrated MHC supertypes can predict Bd susceptibility better than individual MHC alleles, with some supertypes conferring protection and others increasing risk. This suggests that MHC alleles function more as part of a complex network than as independent entities. We also quantified MHC allelic-specific expression and observed individual-level variability in expression across both species. Notably, positive and negative associations among MHC alleles were observed. The similarity of these patterns between the two species suggests the presence of conserved regulatory mechanisms across amphibians. Our results provide valuable resources and insights to advance the understanding of adaptive immunological systems and to develop targeted conservation strategies to preserve biodiversity.

## Introduction

Emerging infectious diseases (EIDs) pose significant threats to global biodiversity, with amphibians being particularly vulnerable. The chytrid fungus, *Batrachochytrium dendrobatidis* (Bd), has caused severe declines and even extirpation of amphibian populations worldwide (Fisher *et al*, 2021; Lips *et al*, 2006; Scheele *et al*, 2019) and extinction of several species is predicted (Alroy, 2015). Bd originated in East Asia (O’Hanlon *et al*, 2018), where some amphibian species may have evolved resistance or tolerance to multiple Bd strains (Fu & Waldman, 2019, 2022). Understanding the genetic basis of resistance in these species is critical for developing effective conservation strategies.

The major histocompatibility complex (MHC) class II (MHC II) plays a pivotal role in the adaptive immune response by binding peptides derived from pathogens and presenting them to T cells (Janeway *et al*, 2001; Unanue *et al*, 2016). MHC II molecules are highly polymorphic, especially in ß1 domain that encode peptide binding sites (PBS), facilitating a wide range of pathogen recognition (Janeway *et al*., 2001; Unanue *et al*., 2016). While a few studies have linked specific MHC II genotypes to resistance to Bd in amphibians (Bataille *et al*, 2015; Cortazar-Chinarro *et al*, 2022; Fu *et al*, 2023a; Kosch *et al*, 2016; Lau *et al*, 2023; Savage & Zamudio, 2011, 2016), no study, to our knowledge, has investigated functional MHC IIß1-based supertypes and associations (*e*.*g*., epistasis) between putative MHC loci.

This study aims to address these gaps by investigating the diversity of MHC II supertypes and allele-expression patterns in an Asian Bd-resistant species (*Bufo gargarizans*) and an Australasian susceptible species (*Litoria caerulea*). The latter species exhibits a mortality rate of about 93.3% from Bd infection (Fu & Waldman, 2019). Using RNA-based genotyping, we previously obtained full-length MHC II β1 sequences from both species (Fu *et al*, 2023b). In this study, we cluster them into functional MHC II supertypes and compare their alpha and beta diversity as well as the allelic expression patterns in the two species with divergent Bd susceptibility. Our results suggest a shift is needed in MHC research in conservation efforts away from simple diversity estimates and towards more complex phenomena such as supertypes and epistatic interactions. This research has important implications for advancing our understanding of differential disease susceptibility and will inform amphibian conservation efforts.

## Results

We modeled five functional supertypes (ST1−ST5) for *B. gargarizans* and *L. caerulea* (Fig. S1c), with detailed allele distributions presented in the amino acid phylogenetic tree (Fig. S2) and alignments (Fig. S3). Both species exhibited four MHC supertypes each with one unique supertype (ST1 for *L. caerulea*; ST4 for *B. gargarizans*) and three shared supertypes (ST2, ST3, and ST5) (Fig. S1d). Most individuals in both species harbored 2 to 3 supertypes (90.9% in *B. gargarizans* and 86.2% in *L. caerulea*) (Fig. 1a).

**Figure 1.**
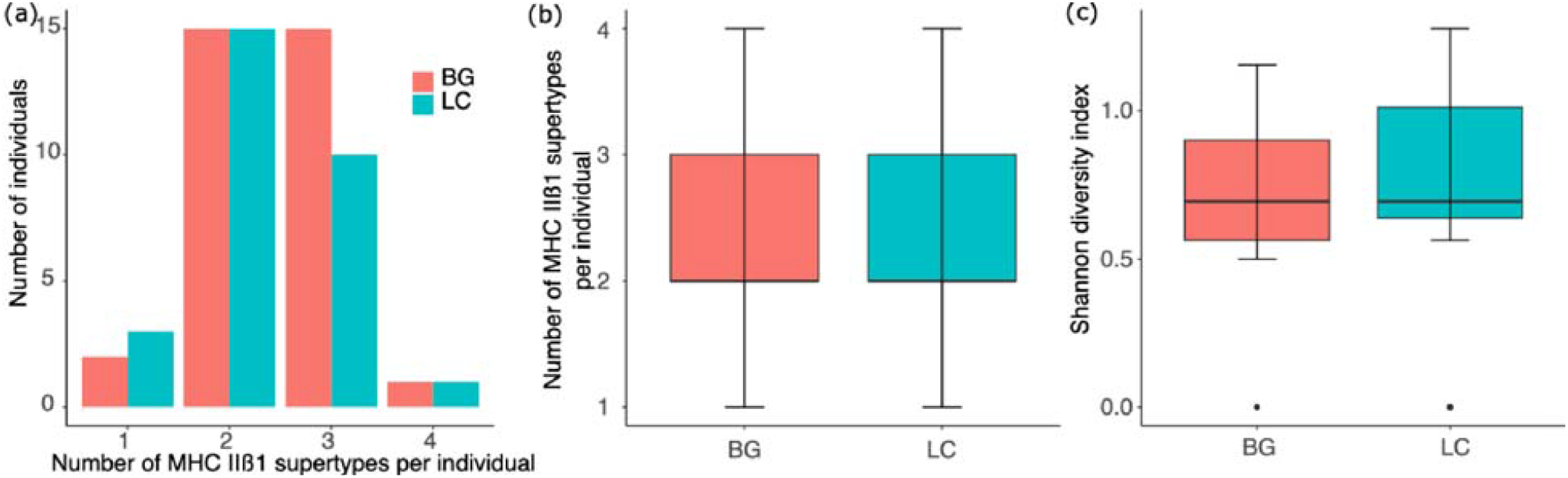
Distribution and alpha diversity of MHC II supertypes in *B. gargarizans* and *L. caerulea*. (a) Distribution of the number of MHC II supertypes per individual in *B. gargarizans* and *L. caerulea*. The majority of individuals in both species possess 2 to 3 supertypes. (b) Comparison of the number of MHC II supertypes between *B. gargarizans* and *L. caerulea*. No significant differences were detected (Wilcoxon rank sum test, *W* = 510.5, *p* = 0.44). (c) Comparison of the Shannon diversity index of MHC II supertypes between *B. gargarizans* and *L. caerulea*. No significant differences were detected (*W* = 389.5, *p* = 0.30). BG, *B. gargarizans*. LC, *L. caerulea*.

### Bd resistance is not associated with MHC supertype alpha diversity

No significant differences were found between the two species in MHC supertype richness (Wilcoxon rank sum test, *W* = 510.5, *p* = 0.44) (Fig. 1b) or Shannon diversity index (*W* = 389.5, *p* = 0.30) (Fig. 1c). Generalized linear mixed effect models (GLMM) indicated that Bd resistance was not associated with alpha diversity metrics (richness, Shannon diversity index, Simpson index) (*p* > 0.05) (Table S1 & S2).

### Bd resistance is associated with MHC supertype components

The compositions of MHC supertypes differed markedly between *B. gargarizans* and *L. caerulea*, with ST5 dominating in the former and ST1 in the latter (Fig. 2a). The structures of the MHC supertype were significantly different between the two species (Fig. 2b), as indicated by PERMANOVA (*F*_1,59_ = 193.84, *p* = 0.0001) and ANOSIM tests (*R* = 0.96, *p* = 0.0001). The GLMMs revealed that ST1 was significantly negatively associated with Bd resistance (Table S1), while ST5 was positively associated (Table S2) (*p* < 0.01). Furthermore, the MHC II supertypes were more dispersed in *L. caerulea* compared to *B. gargarizans* (Betadispersion, *F*_1,59_ = 8.05, *p* = 0.004) (Fig. 2c).

**Figure 2.**
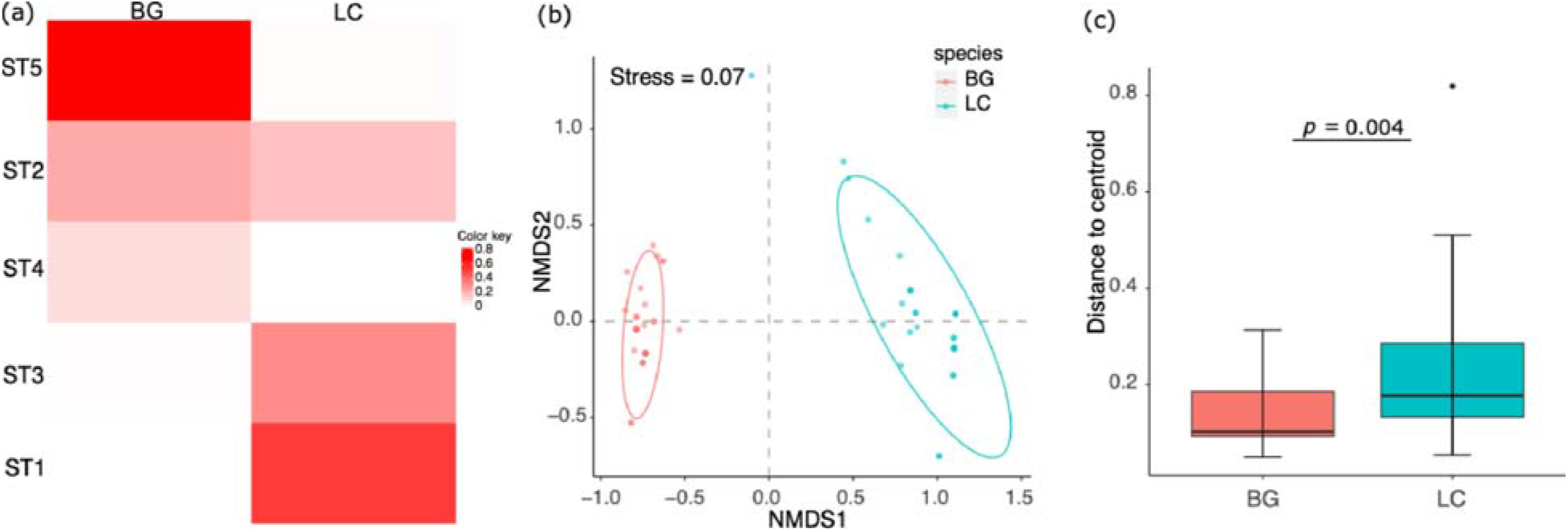
Structure and beta dispersion of MHC II supertypes in *B. gargarizans* and *L. caerulea*. (a) Heatmap plotting relative abundance of MHC II supertypes in both species, ranking from high to low abundance in *B. gargarizan*, with corresponding relative abundance of MHC II supertypes in *L. caerulea*. Unique supertypes for *B. gargarizans* and *L. caerulea* were ST4 and ST1, respectively. (b) Non-metric multidimensional scaling (NMDS) plot showing the distinct structure of MHC II supertypes between *B. gargarizans* and *L. caerulea* (PERMANOVA, *F*_1,59_ = 193.84, *p* = 0.0001). (c) Comparison of beta dispersion for the two species. *L. caerulea* shows more dispersed MHC II supertypes (Betadispersion, *F*_1,59_ = 8.05, *p* = 0.004). Distance to centroid, distance of individual MHC II supertypes to the group centroid. *p* values correspond to PERMDISP (disp) tests. BG, *B. gargarizans*. LC, *L. caerulea*.

### Physicochemical properties of MHC supertypes

The unique supertypes ST1 (*L. caerulea*) and ST4 for (*B. gargarizans*) differed significantly in several physicochemical properties, including Z1 (hydrophobicity) (χ^2^ = 14.08, *p* = 0.0002) (Fig. S4a), Z3 (Polarity) (χ^2^ = 14.08, *p* = 0.0002) (Fig. S4c, Table S5) and Z4 (relating to electronegativity, heat of formation, electrophilicity and hardness) (χ^2^ = 12.62, *p* = 0.0004) (Fig. S4d). Specifically, ST4 exhibited higher hydrophobicity and higher values of Z3 and Z4 compared to ST1 (Fig. S4a). Among shared supertypes (ST2, ST3, and ST5), ST2 showed significantly higher hydrophobicity in *B. gargarizans* compared to *L. caerulea* (Kruskal–Wallis rank sum test, χ^2^ = 8.24, *p* = 0.004) (Fig. S4a). Furthermore, *L. caerulea* exhibited higher Z2 (Steric bulk/Polarizability) for ST2 (χ^2^ = 6.15, *p* = 0.013) and ST5 (χ^2^ = 5.08, *p* = 0.02) than that in *B. gargarizans* (Fig. S4b).

### MHC supertypes predict Bd susceptibility

The relative risk analysis of MHC supertypes in Bd susceptibility revealed significant differences in how various MHC supertypes impact susceptibility to Bd-induced mortality (χ^2^ = 77.39, *df* = 4, *p* < 0.01) (Fig. 3).

**Figure 3.**
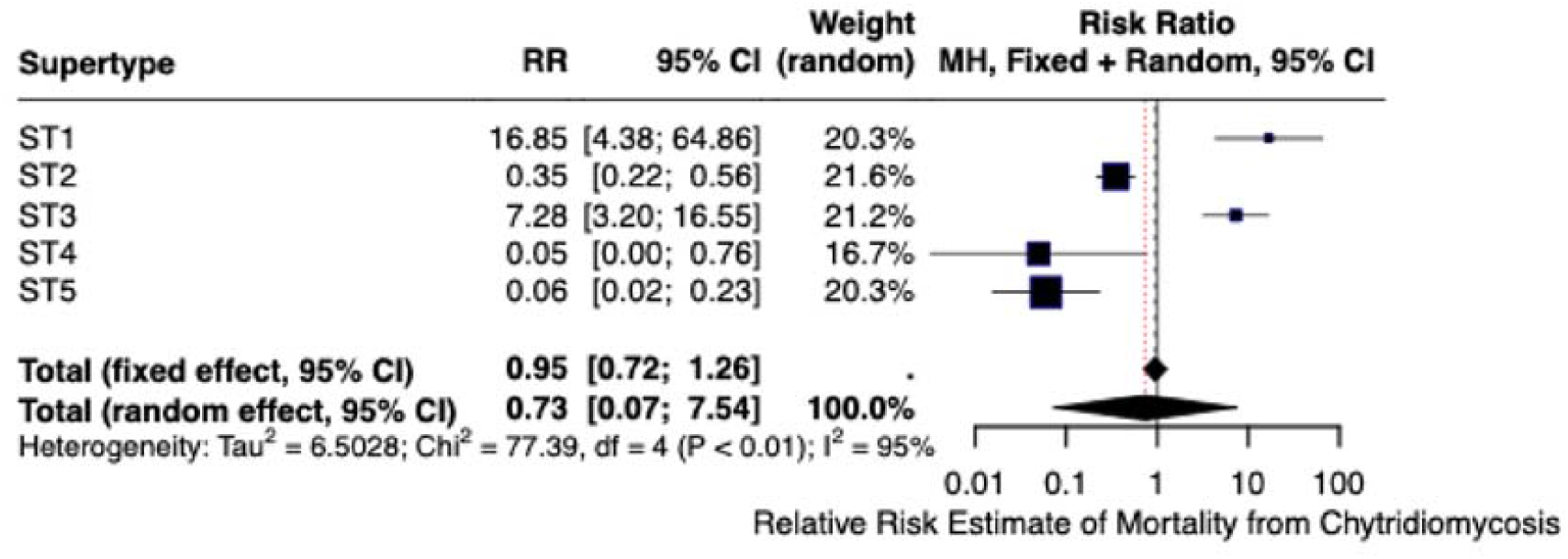
Relative risk estimation of mortality from chytridiomycosis for the MHC supertypes in Bd-resistant and susceptible species.

ST1 and ST3 were identified as high-risk factors for chytridiomycosis-induced mortality, with markedly elevated risk ratios (Fig. 3). Specifically, ST1 had an RR of 16.85 with 95% CI [4.38, 64.86], making it a highly risky supertype linked to Bd susceptibility. Similarly, ST3 also posed a significant risk with an RR of 7.28 [3.20, 16.55] (Fig. 3).

In contrast, ST5, ST4, and ST2 were associated with a protective effect against Bd-induced mortality (Fig. 3). ST5, in particular, demonstrated a strong protective effect with a low RR of 0.06 [0.02, 0.23] (Fig. 3). ST4 and ST2 also exhibited protective roles, with RR of 0.05 [0.00, 0.76] and 0.35 [0.22, 0.56], respectively (Fig. 3).

### MHC alleles fail to predict Bd susceptibility

An individual with Bd-resistant *L. caerulea* (LC15) highly expressed the LitcaeIIB29 allele (Fig. S5), which belongs to the ST2 supertype (Fig. S3b). LitcaeIIB29 contained specific motifs and deletion associated with Bd-resistance (Fu *et al*., 2023a), suggesting it might be a protective allele against Bd infection. However, not all *L. caerulea* individuals expressing LitcaeIIB29 survived Bd infection (that is, LC24, LC1) (Fig. S5), indicating the presence of suppressive alleles for resistance to Bd. Susceptible individuals (e.g., LC6, LC2) expressed other alleles of MHC IIβ1 alleles (e.g., LitcaeIIB14, LitcaeIIB9, LitcaeIIB4, LitcaeIIB26) at high levels (Fig. S5), which belongs mainly to ST1 and ST3 (Fig. S3a &S3c).

A relative risk analysis was performed for these alleles (LitcaeIIB29, LitcaeIIB14, LitcaeIIB9, LitcaeIIB4, LitcaeIIB26), along with frequent alleles (LitcaeIIB19, LitcaeII6) (Fu *et al*., 2023b), and an additional allele in LC15 (LitcaeII24) of *L. caerulea*. The results showed that none of these MHC alleles was significantly associated with Bd susceptibility, with an RR value of approximately 1[0.71, 1.07] (Fig. 4). Furthermore, no significant differences were observed in the risk of susceptibility to Bd between these alleles in *L. caerulea* (χ^2^ = 7.95, *df* = 7, *p* = 0.34) (Fig. 4).

**Figure 4.**
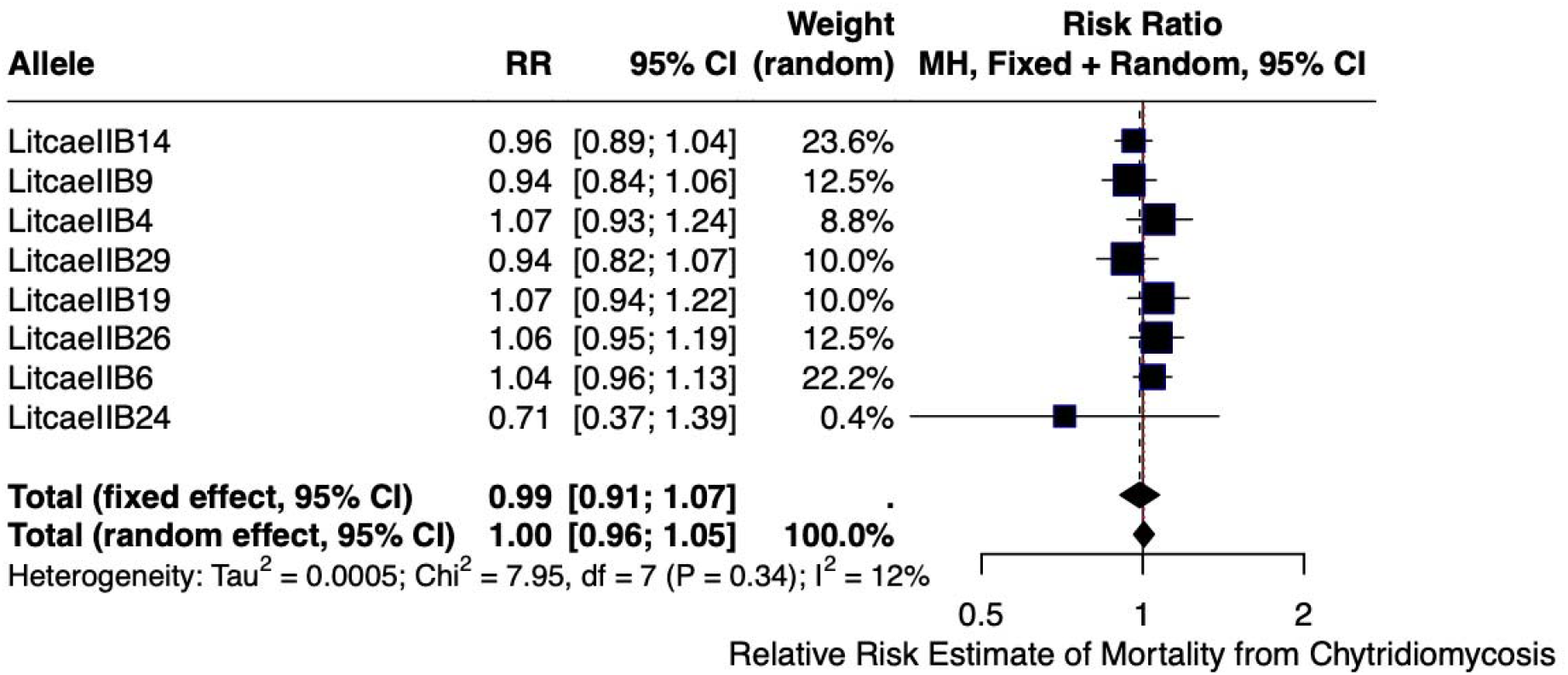
Relative risk estimation of mortality from chytridiomycosis for MHC alleles in Bd-susceptible species, *L. caerulea*.

### Allele-specific expression patterns of MHC IIß1 alleles

As MHC allelic expression varied between some individuals with *L. caerulea* (Fig. S5), we investigated the allelic expression of MHC IIβ1 alleles in both *B. gargarizans* and *L. caerulea*. The expression patterns of the MHC IIß1 alleles in *B. gargarizans* and *L. caerulea* showed consistent variability among individuals, each individual having one or two alleles with high expression levels, while the remaining alleles were expressed at low levels (Fig. 5). Some alleles were expressed in almost all individuals (e.g., BufgarIIB19, BufgarIIB9 in *B. gargarizans*; LitcaeIIB14, LitcaeIIB19 in *L. caerulea*), while other alleles were only expressed in a subset of individuals (e.g., Buga1, Buga3, LC6, LC26) (Figs. 5 & S6–S7).

**Figure 5.**
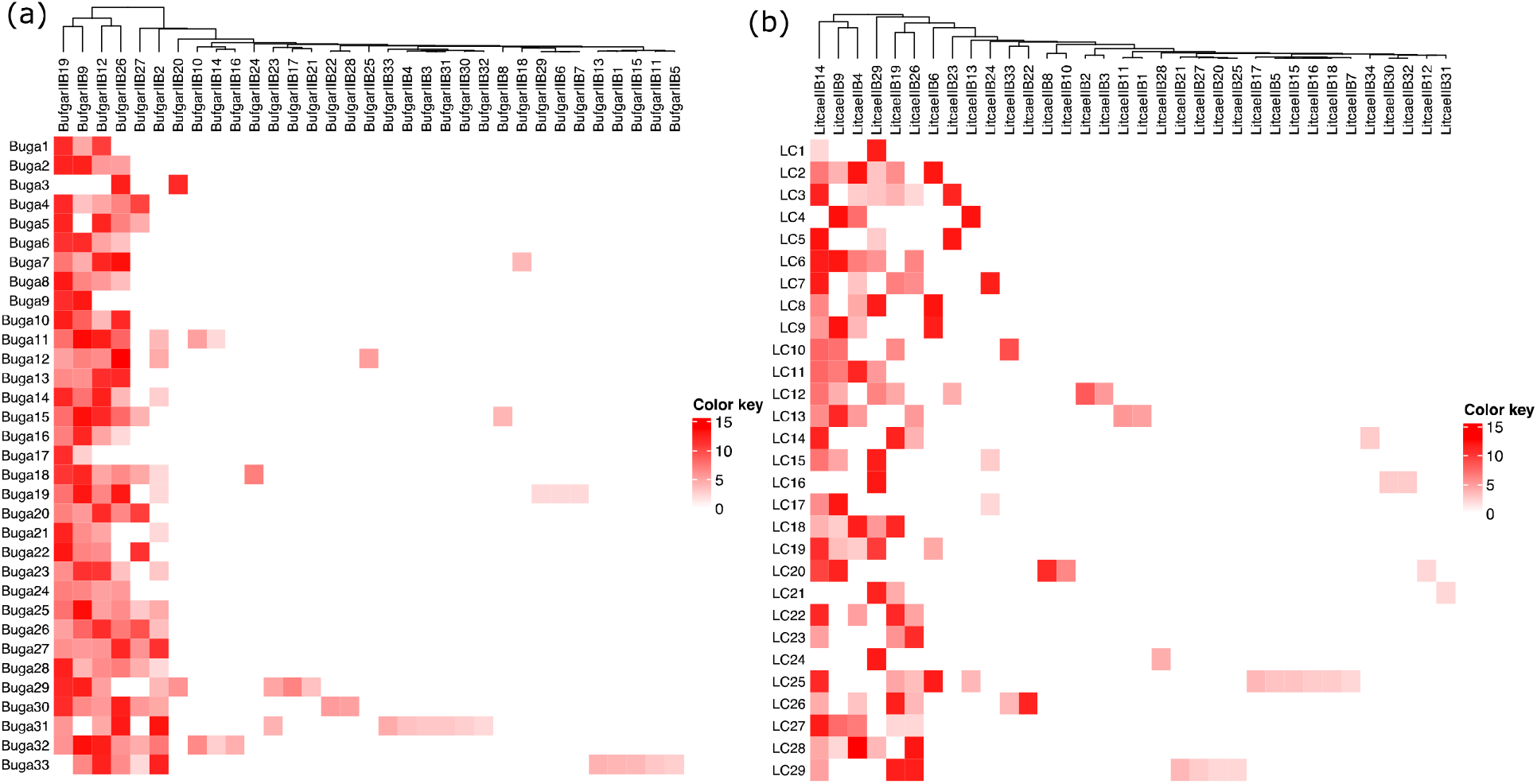
Gene expression patterns of MHC IIβ1 alleles in *B. gargarizans* and *L. caerulea*. Expression levels of MHC IIβ1 alleles in *B. gargarizans* and *L. caerulea*. Each individual typically has one or two highly expressed alleles, with remaining alleles expressed at lower levels. Each lane (such as Buga1) represents an individual. Each individual typically has one or two highly expressed alleles, with remaining alleles expressed at lower levels. Color key represents the log2 normalized transcriptional expression strength, the higher value, the higher expression.

Statistics analysis revealed that there were no significant differences between replicates in beta diversity (PERMANOVA, *B. gargarizans, F*_1,64_ = −0.01, *p* = 0.99; *L. caerulea, F*_1,56_ = 0.03, *p* = 0.99) or in the Shannon diversity index (Wilcoxon rank sum test, *B. gargarizans, W* = 561, *p* = 0.84; *L. caerulea, W* = 429, *p* = 0.90) (Fig. S8).

### Epistatic association among MHC II alleles

Interestingly, when some alleles (e.g., LitcaeIIB29, LitcaeIIB14) were highly expressed, other alleles (e.g., LitcaeIIB14, LitcaeIIB29) were expressed at low levels (Fig. 6b), suggesting a negative correlation between certain MHC II alleles. To explore the intrinsic relationship among all MHC alleles, we performed a Pearson’s correlation analysis and found significant positive and negative associations among MHC II alleles in both species (*B. gargarizans*, Fig. 6a & S9; *L. caerulea*, Fig. 6b &S10). For example, in *B. gargarizans*, BufgarIIB19 was negatively correlated with BufgarIIB26 and BufgarIIB2 significantly (Fig. 6a); in *L. caerulea*, LitcaeIIB29 was negatively correlated with LitcaeIIB14 and LitcaeIIB26 significantly (Fig. 6b). Some alleles exhibited both positive and negative correlations with each other (Fig. 6a & S9: BufgarIIB1, BufgarIIB15, BufgarIIB13, BufgarIIB11, BufgarIIB5, BufgarIIB2; Fig. 6b & S10: LitcaeIIB19,

**Figure 6.**
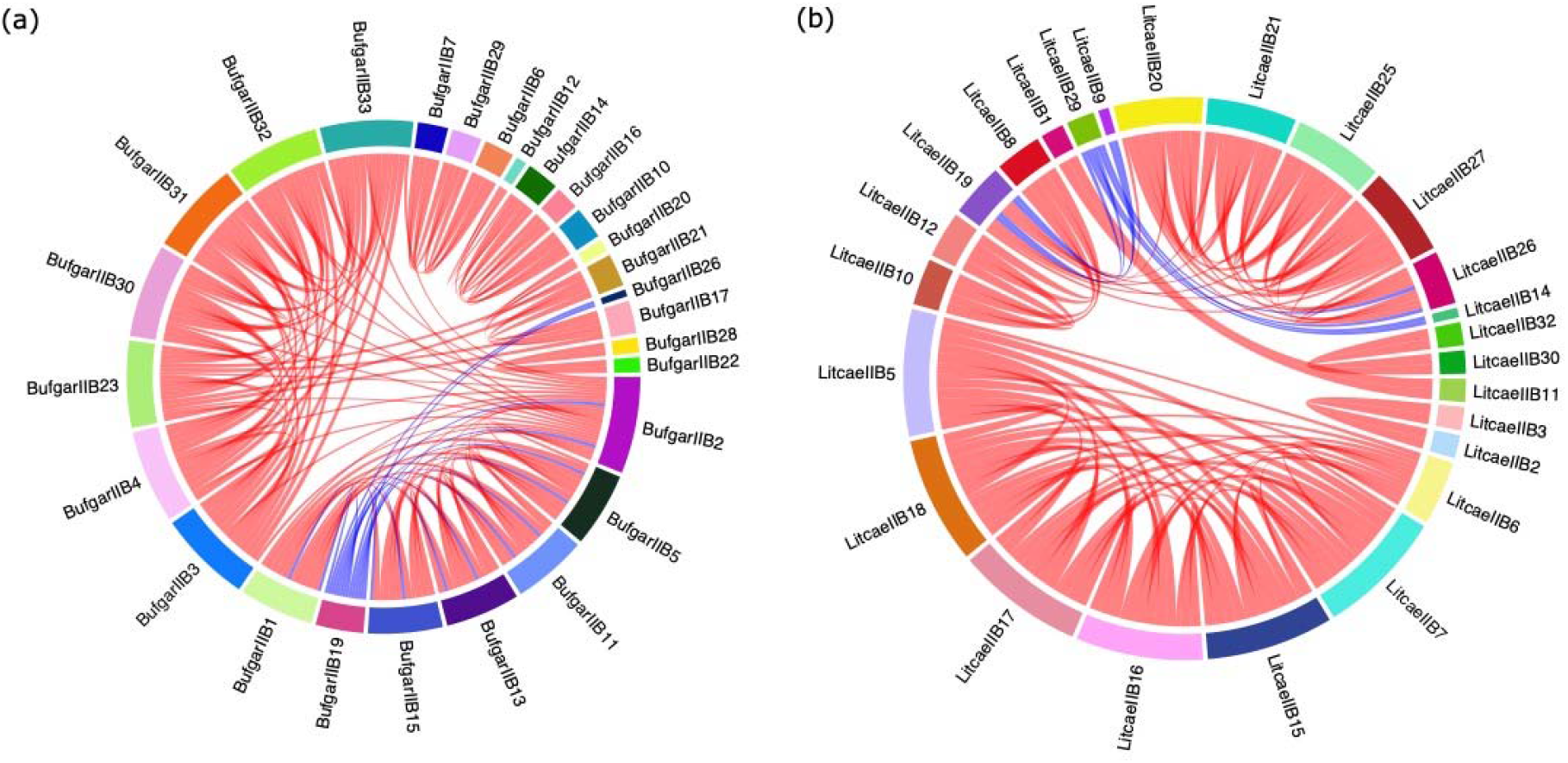
Correlation patterns of MHC II alleles in *B. gargarizans* and *L. caerulea*. (a) Significant (*p* < 0.05) Pearson correlation patterns of MHC II alleles in *B. gargarizans*, showing significant positive and negative associations among alleles (*p* < 0.05). (b) Pearson correlation patterns of MHC II alleles in *L. caerulea*, showing significant positive and negative associations among alleles (*p* < 0.05). Red represents significant positive correlations. Blue represents significant negative correlations.

LitcaeIIB26), but most were positively correlated with each other (Fig. 6). Furthermore, some alleles were exclusively negatively correlated with others (Fig. 6a & S9: BufgarIIB19, BufgarIIB26; Fig. 6b & S10: LitcaeIIB9, LitcaeIIB14, LitcaeIIB29).

## Discussion

Our study applied amphibian-Bd model to understand how the diversity of MHC II functional supertypes contributes to differential Bd susceptibility and their allele-specific expression patterns. Our findings suggest that MHC supertypes, but not MHC alleles, significantly influence risk and protection, thus predicting Bd susceptibility. Transcriptional analysis at the allelic level reveals substantial variability in expression between individuals, along with intrinsic positive and negative correlations between MHC alleles, indicating a complex regulatory network that could shape immune responses across species. These findings deepen our understanding of host immunogenetics and offer valuable information for conservation strategies to mitigate the impact of Bd on vulnerable amphibian populations.

### Bd susceptibility is associated with MHC supertype components rather than alpha diversity

Our modeling reveals both shared and unique supertypes across the two amphibian species (Fig. S1 & Fig. 2). Shared supertypes (ST2, ST3, ST5) can represent conserved immune functions derived from a common ancestor, while unique supertypes (ST1 in *L. caerulea* and ST4 in *B. gargarizans*) can reflect species-specific adaptations to Bd or local environmental pressure as found in fish (Lozano-Martín *et al*, 2023) and bats (Li *et al*, 2023). Our analysis shows that resistance to Bd is not significantly related to the alpha diversity of MHC supertypes or allelic richness (Fig. 1, Table S1), but a significant association exists with beta diversity (Fig. 2). This suggests that Bd susceptibility depends more on the specific supertypes of MHC present than on the overall diversity.

The distinct structural differences in MHC supertypes between *B. gargarizans* and *L. caerulea* also point to an evolutionary divergence in immune system adaptations. The unique supertypes ST1 in *L. caerulea* and ST4 in *B. gargarizans* exhibited significant differences in hydrophobicity, polarity, and other physicochemical properties (Fig. S4). These differences likely underpin the functional divergence between these supertypes, with ST4’s higher hydrophobicity potentially enhancing Bd resistance through improved peptide binding affinity. Such physicochemical properties could be key determinants of MHC-mediated immune responses, as previous studies have also suggested that hydrophobicity selection plays a role in Bd resistance (Fu, 2024).

### MHC II as a double-edged sword in Bd susceptibility

The relative risk (RR) analysis underscores the distinct roles of specific MHC supertypes in resistance to Bd, with ST1 and ST3 identified as risk factors, while ST5, ST4 and ST2 appear to be protective against mortality induced by chytridiomycosis (Fig. 3). These findings reveal that MHC II serve as a double-edged sword in Bd susceptibility, explaining the previous inconsistent findings (Bataille *et al*., 2015; Fu *et al*., 2023a; Savage *et al*, 2020; Savage & Zamudio, 2011, 2016). This dual role underscores the complex nature of MHC-mediated immune responses, where the presence of specific supertypes can increase susceptibility or enhance resistance to pathogens. Understanding this duality is crucial for understanding how MHC diversity influences the outcomes of host-pathogen interactions and has significant implications for the development of conservation strategies to combat chytridiomycosis.

Interestingly, although clear associations between MHC supertype and Bd susceptibility emerged when comparing both Bd-resistant and susceptible species (Fig. 3), no specific MHC alleles in susceptible species exhibited a significant risk of Bd susceptibility (Fig. 4). This suggests that the MHC supertypes predict Bd susceptibility better than alleles. Additionally, it appears that MHC alleles function more as part of a complex network than as independent entities, highlighting the collaborative nature of MHC in modulating disease resistance.

### The intrinsic association among MHC alleles

The multifaced role of MHC is most likely associated with the intrinsic associations among MHC alleles of MHC IIβ1 alleles (Fig. 6), which represents a significant finding of this study. The correlations among the MHC alleles of both species (Fig. 6) suggest potential epistatic regulation, where the expression of one allele can influence others. Epistasis could play a role in synergistically activating positively correlated alleles or antagonistically suppressing others, explaining the inconsistencies in the role of MHC in predicting Bd susceptibility (Fig. 4) and its varied roles.

Epistasis is vital to understanding the complexity of diseases, and is particularly important in polymorphic genes (Mackay & Anholt, 2024; Mackay & Moore, 2014). Further research into the mechanisms behind the specific patterns of MHC alleles and the evolutionary context of potential epistatic interactions could offer valuable insight into how MHC diversity has evolved to optimize resistance to pathogens. Allele-specific expression of MHC should be considered when studying how MHC contributes to differential disease susceptibility. Understanding how the expression of MHC alleles is regulated is essential to advance conservation efforts and disease mitigation strategies.

### Eco-evolutionary significance

The evolutionary basis for these allele-expression patterns and potential epistatic interactions remains an open question. Human MHC (HLA) studies have also found allele-specific expressions (Ramsuran *et al*, 2015; Yamamoto *et al*, 2020), suggesting that this phenomenon for MHC may be conserved across vertebrates, with amphibians providing an evolutionary foundation. Given the key role of MHC in adaptive immunity, its expression throughout the body likely plays a significant role in maintaining homeostasis and health (Janeway *et al*., 2001).

The presence of highly expressed alleles in certain individuals indicates that quantitative differences in gene expression among specific alleles might be crucial to the development of an effective immune response. This can occur synergistically with other expressed MHC alleles, but may also induce autoimmunity or suppress other alleles that could be beneficial in defending against pathogens. It is possible that selective pressures have favored certain allele combinations that maximize immune effectiveness while suppressing non-essential or detrimental responses. The prevalence of positive correlations among MHC alleles than negative correlations (Fig. 6) suggests that while antagonistic effects might have been selected for critical immunoregulatory roles, they may come at a cost.

### Conservation implications

Selective breeding and targeted genetic intervention can offer effective protection for endangered species susceptible to Bd, making it essential to understand the genetic basis of disease susceptibility (Berger *et al*, 2024; Kosch *et al*, 2022). Our results by modelling specific MHC supertypes and conducting risk analysis in Bd-resistant and susceptible species provide evidence that MHC II serves as a double-edged sword for Bd susceptibility. This suggests that targeting the functional MHC supertype may be more effective compared to focusing on a single MHC allele in Bd-susceptible species. These insights lay a critical foundation for targeted conservation efforts, potentially guiding breeding programs or other interventions aimed at bolstering the resilience of vulnerable amphibian populations to Bd. Based on the challenges of our recent research on MHC in Bd resistant and susceptible amphibian species, it is also recommended to breed Bd resistant individuals with others to produce offsprings carrying these alleles. This approach could further our understanding of the role of MHC in Bd resistance and help to preserve rare MHC genotypes, which may also be crucial for the conservation effort.

## Materials and Methods

### Field collection and MHC IIβ1 genotyping

*Bufo gargarizans* (*N* = 33) were collected from Jeonju, Republic of Korea, and *Litoria caerulea* (*N* = 30) were wild-caught in New Guinea (Fu *et al*., 2023b; Fu & Waldman, 2019). These *Bufo gargarizans* were demonstrated to be Bd-resistant, as they rapidly cleared Bd infection, whereas *L. caerulea* were confirmed as a Bd-susceptible species with most succumbing to mortality (Fu & Waldman, 2019). Among the 30 *L. caerulea* individuals, there were two Bd-resistant *L. caerulea* individuals (Fu *et al*., 2023b; Fu & Waldman, 2019). mRNA was first isolated from the liver (*N* = 62) and the buccal swab (*N* = 1) of the two species, amplified in duplicate or triplicate, and sequenced using the Illumina MiSeq 2 × 300 bp paired end (PE) Illumina MiSeq platform (Fu *et al*., 2023b). MHC IIβ1 alleles were cloned and sequenced in a previous study (Fu *et al*., 2023b). Thirty-three alleles were identified in *B. gargarizans* and 35 in *L. caerulea* (GenBank accession numbers OQ726420–OQ726487).

### Supertyping for MHC IIβ1 alleles

To cluster MHC IIβ1 into different functional supertypes, we first extracted 24 PBSs from MHC IIβ1 amino acid alignments (*N* = 68) of *B. gargarizans* (*N* = 33) and *L. caerulea* (*N* = 35) (site 9, 11, 13, 28, 30, 32, 37, 38, 47, 56, 60, 61, 65, 68, 70, 71, 74, 78, 81, 82, 85, 86, 88, 89) (Fu *et al*., 2023b). These PBSs were annotated by comparing them with those of the MHC II of *Homo sapiens* (Bergström *et al*, 1998). Each amino acid residue was then characterized based on five physicochemical descriptor variables (Z1, lipophilicity/hydrophobicity; Z2, polarization/steric bulk; Z3, polarity/charge; Z4 and Z5, relate to electronegativity, heat of formation, electrophilicity and hardness) (Sandberg *et al*, 1998).

Next, a discriminant analysis of the principal components (DAPC) was performed to group functionally similar MHC II alleles into different clusters using the adegenet 2.1.10 package in R v. 4.2.1 (Jombart, 2008; Jombart *et al*, 2010). For DAPC, clustering of K-means with Bayesian information criterion (BIC) was used to evaluate the optimal number of clusters (BIC was approximately 60) (Fig. S1a), and cross-validation was used to select the number of principal components (PCs) to be objectively retained. The membership probability for each MHC II allele was shown in Fig. S1b.

### Construction of a phylogenetic tree

To visualize the distribution of MHC II supertypes in *L. caerulea* and *B. gargarizans* in the amino acid phylogenetic tree, we constructed a consensus maximum likelihood tree of full-length MHC IIβ1 alleles with 1000 bootstrap replicas using MEGA X with the JTT + G + I substitution model (Kumar *et al*, 2018). All positions with < 95% site coverage were eliminated; that is, fewer than 5% alignment gaps, missing data, and ambiguous bases were allowed at any position (partial deletion option).

### Alpha and Beta diversity analysis and corresponding statistics

To understand whether resistance to Bd in the two species is related to the alpha diversity of MHC II supertypes, we calculate the number of MHC supertypes (richness), Shannon’s diversity index, Simpson’s index for each individual in Bd-resistant species, *B. gargarizans* (*N* = 33) and Bd-susceptible *L. caerulea* (*N* = 28), using the ‘diversity’ function ‘diversity’ from vegan package (Oksanen, 2013). Two Bd-resistant *L. caerulea* individuals were removed for this comparison with *B. gargarizans* to avoid bias from individuals resistant to Bd from *L. caerulea*. Kruskal–Wallis rank sum tests were conducted using ‘kruskal.test’ in R 3.3.1 (R Foundation for Statistical Computing, Vienna, Austria). Wilcoxon rank sum tests were computed using ‘wilcox.test’ from package ‘MASS’ (Ripley *et al*, 2013).

To determine whether the difference between MHC II supertype components among individuals (beta diversity) can be detected between Bd resistant and susceptible species, nonmetric multidimensional scaling (NMDS), as a multivariate statistical ordination technique (Pierini *et al*, 2018), was used to display the structure of the MHC II supertypes in the two species, based on the relative abundance of each supertype at the individual level. Permutational Multivariate Analysis of Variance (PERMANOVA) and Analysis of Similarities (ANOSIM) were then used to test the statistical differences and similarities between the structure of MHC II supertype components in the Bd-susceptible and resistant species, respectively. PERMANOVA and ANOSIM were based on Bray-Curtis dissimilarity with 9999 permutations using the vegan package (Oksanen, 2013).

To test whether beta dispersion differs between Bd resistant and susceptible species, a beta dispersion test was performed using the ‘betadisper’ function in the vegan package, followed by the permutation test with 9999 permutations using ‘permutest’ in the vegan package to determine statistical significance.

### Assessing the effect of MHC II β1 diversity on Bd resistance

The diversity data of the MHC II supertypes (richness, Shannon’s diversity index, Simpson’s index, beta dispersion, relative abundance of each supertype) generated above for each individual from Bd-resistant *B. gargarizans* and susceptible *L. caerulea* were further used to evaluate whether these factors of the MHC II supertypes were related to resistance to Bd using generalized linear mixed effects models (GLMM) with the function ‘glmer’ in the package of lmer4 (Bates *et al*, 2014). In addition, number of MHC II alleles at the individual level from a previous study (Fu *et al*., 2023b) was also evaluated for Bd resistance using GLMM. The GLMMs were fitted with binary Bd susceptibility data from both species (*N* = 61) as dependent response, MHC IIβ1 diversity as fixed variable, individual ID and species ID as random variables, and fitted with binomial error distribution. Each diversity indices was modeled separately as they were proved to have autocorrelation.

An individual resistant to Bd (LC15) was added (*N* = 62). The beta dispersion was recalculated using the same method as above. The GLMMs were fitted with binary Bd susceptibility data from both species (*N* = 62) as dependent response, MHC IIβ1 diversity as fixed variable, individual ID and species ID as random variables, and fitted with binomial error distribution.

### Physicochemical analysis for MHC IIβ1 supertypes

To understand whether different functional supertypes vary in the five Z variables with respect to Bd resistance in the Bd-resistant species (*B. gargarizans*) and susceptible species (*L. caerulea*), we performed comparisons of the composite values Z among different supertypes using Kruskal-Wallis rank sum tests. A large negative value of Z1 refers to a lipophilic/hydrophobic amino acid residue, a large positive value of Z1 refers to a polar hydrophilic amino acid residue (Sandberg *et al*., 1998). As the LitcaeIIB35 allele was a unique allele of a survivor of *L. caerulea*, it was removed from this analysis.

For comparing the physicochemical properties of particular amino acid residues, Kruskal–Wallis rank sum tests were conducted using the kruskal.test in R.

### Relative risk analysis

Binary data (presence or absence of certain MHC supertypes) were collected from *B. gargarizans* (*N* = 33) and *L. caerulea* (*N* = 29) (Note: One Bd-resistant individual from *L. caerulea* was excluded to avoid tissue bias, as its MHC genotype was obtained from a buccal swab rather than the liver, unlike the other individuals) to evaluate the effect of MHC II supertypes on the mortality risk of chytridiomycosis. The number of events (mortality) and the total number of certain MHC supertypes were collected for each individual for relative risk estimation.

For relative risk analysis in *L. caerulea*, binary data (presence or absence of certain MHC alleles) were collected from *L. caerulea* to assess the effect of MHC II alleles on the mortality risk of chytridiomycosis. A *L. caerulea* individual survived from Bd infection (Fu & Waldman, 2019) and was recorded as 0. Other individuals of *L. caerulea* who succumbed to the Bd infection (Fu & Waldman, 2019) were recorded as 1. The number of mortality events and the total number of individual MHC alleles were collected for each individual to estimate relative risk. Particular MHC alleles were selected for this analysis, with further details provided in the Results section.

Relative risk estimation was performed using the ‘metabin’ function ‘metabin’ from the meta package (Schwarzer, 2007) (version 7.0-0) in R 4.2.1. Fixed and random-effect models were used to estimate the pooled relative risk (RR) of MHC II supertypes and alleles on the mortality risk of chytridiomycosis. When the value of the relative risk (RR) is equal to 1, it indicates that there is no effect. An RR greater than 1 suggests a positive correlation with risk, and we consider it a significant positive correlation when the RR exceeds 2. On the contrary, an RR less than 1 indicates a negative correlation with risk, and we consider it significantly negative when the RR is below 0.5.

### MHC allele-specific expression analysis

To further understand the complexity of MHC, we analyzed the allele-specific transcriptional expression of MHC II. As the sequences were derived from mRNA, it was assumed that the read counts reflect the relative gene expression of the MHC II alleles. Read count data for MHC were obtained from the supplementary material of two previous studies (Fu *et al*., 2023a; Fu *et al*., 2023b). The read counts for the MHC II alleles were normalized using a log transformation on the base of 2 (log_2_Reads). The lengths of all MHC II sequences were not considered for this gene expression analysis in this study, as these MHC II sequences for each species only have 3 bp differences in length (Fu *et al*., 2023a; Fu *et al*., 2023b). Next, heat maps using hierarchical cluster analysis (Murtagh & Legendre, 2014) were used to visualize the relative gene expressions of MHC II alleles for each individual with replicates (Individual_1, Individual_2, and/or Individual_3) in Bd resistant (*B. gargarizans*) and susceptible species (*L. caerulea*).

Replicate differences among Individuals were compared for both species. The beta diversity and Shannon diversity index were plotted to compare whether there were differences between replicates. PERMANOVA based on Bray-Curtis dissimilarity with 9999 permutations using the vegan package was used to compare beta diversity (Oksanen, 2013). The Wilcoxon rank sum test with continuity correction was performed to compare the differences in Shannon diversity index between replicates.

### Correlation analysis

To detect whether there was a correlation in the relative gene expression of MHC II alleles at the individual level, a pairwise Pearson correlation analysis was performed using the relative gene expression of MHC II for each individual with the ‘cor’ function in R. For chord-gram plotting, only significant correlations (*p* < 0.05) were used.

All statistical tests were conducted in R v. 4.2.1 (The R Development Core Team, 2022).

## Acknowledgement

The research was supported by the grant awarded to MF by National Research Foundation of Korea (NRF), funded by the Ministry of Education (2022R1I1A1A01072313) of the Republic of Korea. We would like to thank Prof. Rajendra Karki from Seoul National University for facilitating completion of this research by hosting the grant.

## Conflicts of interest

The authors declare no conflicts of interest.

## Notes

### Competing Interest Statement

The authors have declared no competing interest.

### Summary of Updates

Typos in abstract, methods, and figures are revised.

## References

Alroy J (2015) Current extinction rates of reptiles and amphibians. Proceedings of the National Academy of Sciences 112: 13003–13008

Bataille A, Cashins SD, Grogan L, Skerratt LF, Hunter D, McFadden M, Scheele B, Brannelly LA, Macris A, Harlow PS (2015) Susceptibility of amphibians to chytridiomycosis is associated with MHC class II conformation. Proc R Soc B 282: 20143127

Bates D, Mächler M, Bolker B, Walker S (2014) Fitting linear mixed-effects models using lme4. arXiv preprint arXiv:14065823

Berger L, Skerratt LF, Kosch TA, Brannelly LA, Webb RJ, Waddle AW (2024) Advances in Managing Chytridiomycosis for Australian Frogs: Gradarius Firmus Victoria. Annual Review of Animal Biosciences 12: 113–133

Bergström TF, Josefsson A, Erlich HA, Gyllensten U (1998) Recent origin of HLA-DRB1 alleles and implications for human evolution. Nature Genet 18: 237–242

Cortazar-Chinarro M, Meurling S, Schroyens L, Siljestam M, Richter-Boix A, Laurila A, Höglund J (2022) Major histocompatibility complex variation and haplotype associated survival in response to experimental infection of two Bd-GPL strains along a latitudinal gradient. Frontiers in Ecology and Evolution 10: 915271

Fisher MC, Pasmans F, Martel A (2021) Virulence and Pathogenicity of Chytrid Fungi Causing Amphibian Extinctions. Annual review of microbiology 75: 673–693

Fu M (2024) Evolutionary analysis of major histocompatibility complex variants in chytrid-resistant and susceptible amphibians. Infection, Genetics and Evolution 118: 105544

Fu M, Eimes JA, Kong S, Lamichhaney S, Waldman B (2023a) Identification of major histocompatibility complex genotypes associated with resistance to an amphibian emerging infectious disease. Infection, Genetics and Evolution: 105470

Fu M, Eimes JA, Waldman B (2023b) Divergent allele advantage in the MHC and amphibian emerging infectious disease. Infection, Genetics and Evolution: 105429

Fu M, Waldman B (2019) Ancestral chytrid pathogen remains hypervirulent following its long coevolution with amphibian hosts. Proc R Soc B 286: 20190833

Fu M, Waldman B (2022) Novel chytrid pathogen variants and the global amphibian pet trade. Conservation Biology

Janeway C, Charles A, Travers P, Walport M, Shlomchik MJ (2001) Principles of innate and adaptive immunity. In: Immunobiology: The Immune System in Health and Disease 5th edition, Garland Science:

Jombart T (2008) adegenet: a R package for the multivariate analysis of genetic markers. Bioinformatics 24: 1403–1405

Jombart T, Devillard S, Balloux F (2010) Discriminant analysis of principal components: a new method for the analysis of genetically structured populations. BMC Genetics 11: 94

Kosch TA, Bataille A, Didinger C, Eimes JA, Rodríguez-Brenes S, Ryan MJ, Waldman B (2016) Major histocompatibility complex selection dynamics in pathogen-infected túngara frog (Physalaemus pustulosus) populations. Biol Lett 12: 20160345

Kosch TA, Waddle AW, Cooper CA, Zenger KR, Garrick DJ, Berger L, Skerratt LF (2022) Genetic approaches for increasing fitness in endangered species. Trends in Ecology & Evolution 37: 332–345

Kumar S, Stecher G, Li M, Knyaz C, Tamura K (2018) MEGA X: Molecular Evolutionary Genetics Analysis across Computing Platforms. Mol Biol Evol 35: 1547–1549

Lau Q, Igawa T, Kosch TA, Dharmayanthi AB, Berger L, Skerratt LF, Satta Y (2023) Conserved Evolution of MHC Supertypes among Japanese Frogs Suggests Selection for Bd Resistance. Animals (Basel) 13

Li X, Liu T, Li A, Xiao Y, Sun K, Feng J (2023) Diversifying selection and climatic effects on major histocompatibility complex class II gene diversity in the greater horseshoe bat. Evolutionary Applications 16: 688–704

Lips KR, Brem F, Brenes R, Reeve JD, Alford RA, Voyles J, Carey C, Livo L, Pessier AP, Collins JP (2006) Emerging infectious disease and the loss of biodiversity in a Neotropical amphibian community. Proc Natl Acad Sci USA 103: 3165–3170

Lozano-Martín C, Bracamonte SE, Barluenga M (2023) Evolution of MHC IIB Diversity Across Cichlid Fish Radiations. Genome Biology and Evolution 15: evad110

Mackay TFC, Anholt RRH (2024) Pleiotropy, epistasis and the genetic architecture of quantitative traits. Nature Reviews Genetics 25: 639–657

Mackay TFC, Moore JH (2014) Why epistasis is important for tackling complex human disease genetics. Genome Medicine 6: 42

Murtagh F, Legendre P (2014) Ward’s Hierarchical Agglomerative Clustering Method: Which Algorithms Implement Ward’s Criterion? Journal of Classification 31: 274–295

O’Hanlon SJ, Rieux A, Farrer RA, Rosa GM, Waldman B, Bataille A, Kosch TA, Murray KA, Brankovics B, Fumagalli M (2018) Recent Asian origin of chytrid fungi causing global amphibian declines. Science 360: 621–627

Oksanen J (2013) Vegan: ecological diversity. R Project 368

Pierini F, Lenz TL, Wilke C (2018) Divergent allele advantage at human MHC genes: signatures of past and ongoing selection. Mol Biol Evol

Ramsuran V, Kulkarni S, O’Huigin C, Yuki Y, Augusto DG, Gao X, Carrington M (2015) Epigenetic regulation of differential HLA-A allelic expression levels. Hum Mol Genet 24: 4268–4275

Ripley B, Venables B, Bates DM, Hornik K, Gebhardt A, Firth D, Ripley MB (2013) Package ‘mass’. Cran r 538: 113–120

Sandberg M, Eriksson L, Jonsson J, Sjöström M, Wold S (1998) New chemical descriptors relevant for the design of biologically active peptides. A multivariate characterization of 87 amino acids. J Med Chem 41: 2481–2491

Savage AE, Gratwicke B, Hope K, Bronikowski E, Fleischer RC (2020) Sustained immune activation is associated with susceptibility to the amphibian chytrid fungus. Mol Ecol 29: 2889–2903

Savage AE, Zamudio KR (2011) MHC genotypes associate with resistance to a frog-killing fungus. Proc Natl Acad Sci U S A 108: 16705–16710

Savage AE, Zamudio KR (2016) Adaptive tolerance to a pathogenic fungus drives major histocompatibility complex evolution in natural amphibian populations. Proc R Soc B 283: 20153115

Scheele BC, Pasmans F, Skerratt LF, Berger L, Martel A, Beukema W, Acevedo AA, Burrowes PA, Carvalho T, Catenazzi A (2019) Amphibian fungal panzootic causes catastrophic and ongoing loss of biodiversity. Science 363: 1459–1463

Schwarzer G (2007) meta: An R package for meta-analysis. R news 7: 40–45

Unanue ER, Turk V, Neefjes J (2016) Variations in MHC class II antigen processing and presentation in health and disease. Annu Rev Immunol 34: 265–297

Yamamoto F, Suzuki S, Mizutani A, Shigenari A, Ito S, Kametani Y, Kato S, Fernandez-Viña M, Murata M, Morishima S et al (2020) Capturing Differential Allele-Level Expression and Genotypes of All Classical HLA Loci and Haplotypes by a New Capture RNA-Seq Method. Frontiers in Immunology 11

