## Supplementary for "The major histocompatibility complex: a double-edged sword in fungal disease susceptibility"

Table S1 Generalized linear mixed-effects models (GLMMs) for the effects of alpha diversity indices and the relative abundance of each MHC supertypes on Bd Resistance.

|  | Estimate | Std. Error | *z* | *Pr(>\|z\|)* | mR^2^ | cR^2^ |
| --- | --- | --- | --- | --- | --- | --- |
| (Intercept) | 14.98 | 14.32 | 1.05 | 0.30 | 0.01 | 1.00 |
| richness of supertypes | 0.14 | 5.68 | 0.03 | 0.98 |  |  |
| (Intercept) | 15.42 | 11.35 | 1.36 | 0.17 | 0.00 | 1.00 |
| Shannon of supertypes | -0.13 | 13.01 | -0.01 | 0.99 |  |  |
| (Intercept) | 15.58 | 9.97 | 1.56 | 0.12 | 0.01 | 1.00 |
| Simpson of supertypes | -0.58 | 17.15 | -0.03 | 0.97 |  |  |
| (Intercept) | 15.92 | 8.44 | 1.89 | 0.06 | 0.13 | 1.00 |
| beta dispersion of supertypes | -4.05 | 24.57 | -0.17 | 0.87 |  |  |
| (Intercept) | 14.83 | 11.46 | 1.30 | 0.20 | 0.03 | 1.00 |
| richness of alleles | 0.09 | 2.05 | 0.05 | 0.96 |  |  |
| (Intercept) | 14.58 | 8.73 | 1.67 | 0.09 | 0.88 | 1.00 |
| **ST1** | **-5624.85** | **1024.00** | **-5.49** | **0.00** |  |  |
| (Intercept) | 15.14 | 7.50 | 2.02 | 0.04 | 0.04 | 1.00 |
| ST2 | 0.82 | 15.40 | 0.05 | 0.96 |  |  |
| (Intercept) | 15.07 | 6.63 | 2.27 | 0.02 | 0.68 | 1.00 |
| ST3 | -12.42 | 21.96 | -0.57 | 0.57 |  |  |
| (Intercept) | 14.50 | 79.07 | 0.18 | 0.85 | 0.30 | 1.00 |
| ST4 | 227.50 | 82440000.00 | 0.00 | 1.00 |  |  |
| (Intercept) | -94.66 | 12570000.00 | 0.00 | 1.00 | 1.00 | 1.00 |
| ST5 | 351.10 | 24820000.00 | 0.00 | 1.00 |  |  |

Alpha diversity indices: richness, Shannon’s diversity index, Simpson’s index. mR^2^, marginal R^2^, explaining the variance of the fixed effect in the model. cR^2^, conditional R^2^, explaining the variance of both the fixed and random effects. *N* = 61.

Table S2 Generalized linear mixed-effects models (GLMMs) for the effects of alpha diversity indices and the relative abundance of each MHC supertypes on Bd Resistance.

|  | Estimate | Std. Error | *z* | *Pr(>\|z\|)* | mR^2^ | cR^2^ |
| --- | --- | --- | --- | --- | --- | --- |
| (Intercept) | 11.05 | 8.19 | 1.35 | 0.18 | 0.01 | 0.94 |
| richness of supertypes | -0.58 | 1.43 | -0.40 | 0.69 |  |  |
| (Intercept) | 10.57 | 7.44 | 1.42 | 0.16 | 0.00 | 0.94 |
| Shannon of supertypes | -1.40 | 2.76 | -0.51 | 0.61 |  |  |
| (Intercept) | 10.50 | 7.42 | 1.42 | 0.16 | 0.01 | 0.94 |
| Simpson of supertypes | -2.23 | 4.38 | -0.51 | 0.61 |  |  |
| (Intercept) | 9.34 | 6.99 | 1.34 | 0.18 | 0.12 | 0.94 |
| beta dispersion of supertypes | 1.35 | 5.27 | 0.26 | 0.80 |  |  |
| (Intercept) | 10.60 | 8.25 | 1.28 | 0.20 | 0.03 | 0.94 |
| richness of alleles | -0.18 | 0.61 | -0.29 | 0.77 |  |  |
| (Intercept) | 10.48 | 6.47 | 1.62 | 0.11 | 0.78 | 0.94 |
| ST1 | 4.10 | 5.10 | 0.80 | 0.42 |  |  |
| (Intercept) | 9.15 | 7.13 | 1.28 | 0.20 | 0.04 | 0.94 |
| ST2 | 1.26 | 3.42 | 0.37 | 0.71 |  |  |
| (Intercept) | 4.06 | 8.18 | 0.50 | 0.62 | 0.68 | 0.94 |
| ST3 | -17.98 | 20.48 | -0.88 | 0.38 |  |  |
| (Intercept) | 8.34 | 9.87 | 0.85 | 0.40 | 0.29 | 0.94 |
| ST4 | 214.70 | 64920000.00 | 0.00 | 1.00 |  |  |
| (Intercept) | -3.66 | 1.12 | -3.27 | 0.00 | 0.93 | 0.93 |
| **ST5** | **16.30** | **5.85** | **2.79** | **0.01** |  |  |

Alpha diversity indices: richness, Shannon’s index, Simpson’s index. mR^2^, marginal R^2^, explaining the variance of the fixed effect in the model. cR^2^, conditional R^2^, explaining the variance of both the fixed and random effects. *N* = 62.

Table S3 Statistics from Kruskal–Wallis rank sum tests for physicochemical analysis of MHC II supertypes in Bd-resistant (*B. gargarizans*) and susceptible species (*L. caerulea*).

|  | Groups | *c^2^* | *df* | *p* |
| --- | --- | --- | --- | --- |
| **Z1** | **ST2(LC), ST2(BG)** | **8.25** | **1.00** | **0.00** |
|  | ST3(LC), ST3(BG) | 1.80 | 1.00 | 0.18 |
|  | ST5(LC), ST5(BG) | 0.71 | 1.00 | 0.40 |
|  | **ST1(LC), ST4(BG)** | **14.08** | **1.00** | **0.00** |
| **Z2** | **ST2(LC), ST2(BG)** | **6.15** | **1.00** | **0.01** |
|  | ST3(LC), ST3(BG) | 0.07 | 1.00 | 0.79 |
|  | **ST5(LC), ST5(BG)** | **5.08** | **1.00** | **0.02** |
|  | ST1(LC), ST4(BG) | 0.54 | 1.00 | 0.46 |
| **Z3** | ST2(LC), ST2(BG) | 0.04 | 1.00 | 0.84 |
|  | ST3(LC), ST3(BG) | 1.15 | 1.00 | 0.28 |
|  | ST5(LC), ST5(BG) | 0.08 | 1.00 | 0.78 |
|  | **ST1(LC), ST4(BG)** | **14.08** | **1.00** | **0.00** |
| **Z4** | ST2(LC), ST2(BG) | 2.79 | 1.00 | 0.09 |
|  | ST3(LC), ST3(BG) | 0.65 | 1.00 | 0.42 |
|  | ST5(LC), ST5(BG) | 0.71 | 1.00 | 0.40 |
|  | **ST1(LC), ST4(BG)** | **12.62** | **1.00** | **0.00** |
| Z5 | ST2(LC), ST2(BG) | 1.29 | 1.00 | 0.26 |
|  | ST3(LC), ST3(BG) | 2.59 | 1.00 | 0.11 |
|  | ST5(LC), ST5(BG) | 2.86 | 1.00 | 0.09 |
|  | ST1(LC), ST4(BG) | 0.54 | 1.00 | 0.46 |

BG, *B. gargarizans*. LC, *L. caerulea*

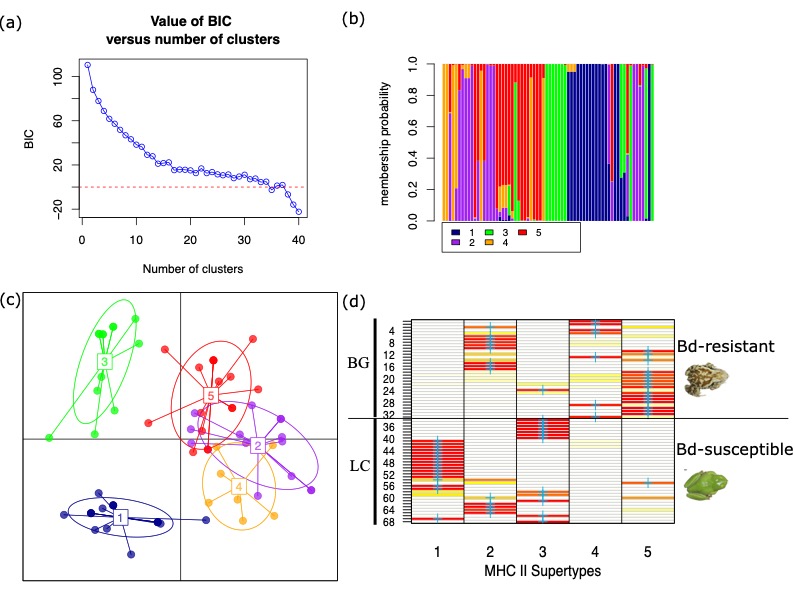

Figure S1 Characterization of MHC II supertypes in Bd-resistant and susceptible species. (a) Bayesian information criterion (BIC) values relative to the number of clusters. (b) **Membership probability of MHC IIβ1 alleles in predicted supertypes.** The bar plot displays the membership probability of each MHC IIβ1 allele to the five predicted supertypes. Each bar represents an allele, and the colored segments within each bar indicate the probability of belonging to each supertype. (c) **Discriminant Analysis of Principal Components (DAPC)** plot shows the clustering of MHC IIβ1 alleles into five distinct supertypes for Bufo gargarizans and Litoria caerulea. Each point represents an allele, with different colors indicating different supertypes. The axes represent the first two principal components, which capture the most variation in the dataset. Each number represent a predicted supertype. (d) distribution of MHC II supertypes in Bd-resistant and susceptible species. The first 33 lanes represent 33 MHC II alleles from Bd-resistant species, *B. gargarizans* (BG). The last 35 lanes represents 35 MHC II alleles from Bd-susceptible species, *L. caerulea* (LC).

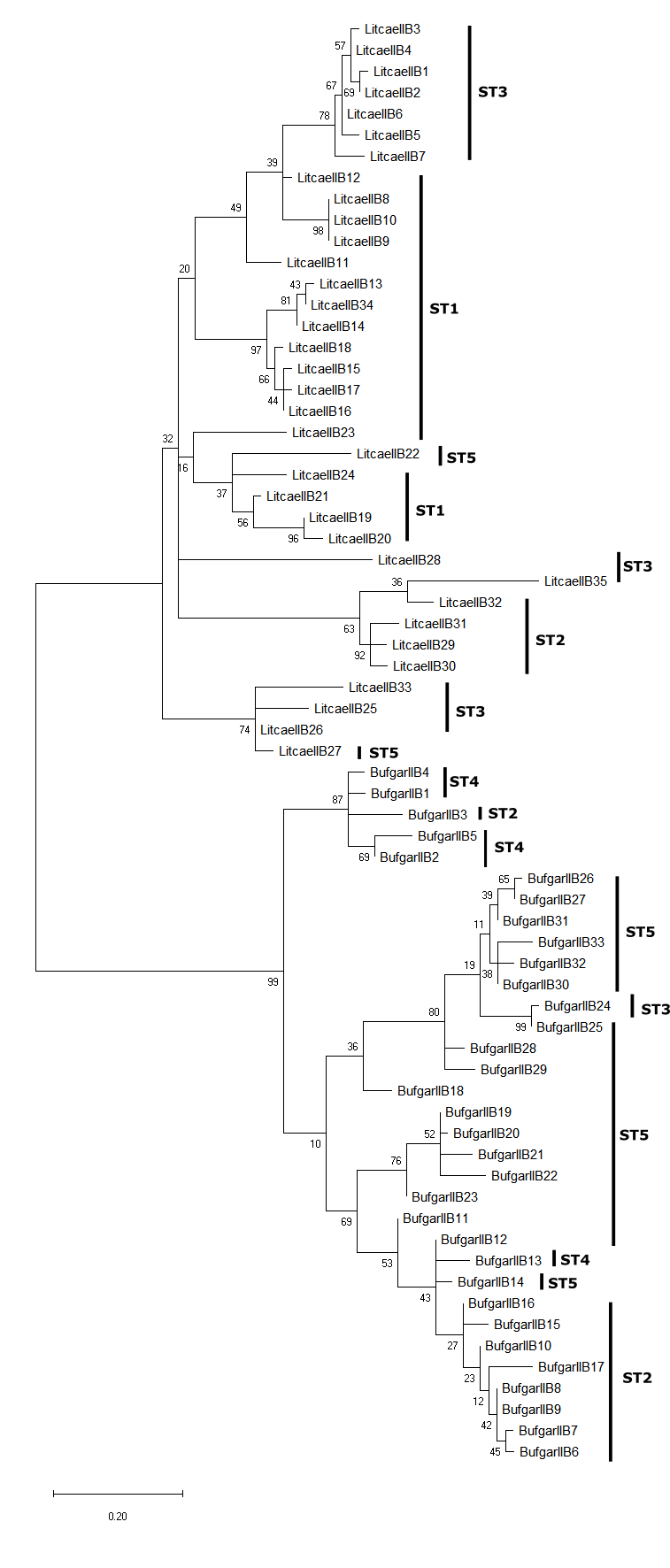

Figure S2 Distribution of MHC II supertypes in the full MHC IIß1 amino acid sequences-based maximum likelihood phylogenetic tree for MHC II alleles in Bd-resistant and susceptible species. Branch lengths were measured in the number of substitutions per site. One thousand bootstrap replicates were conducted. The values next to branches represent bootstrap values. Note: MHC IIB1 is synonymous with MHC IIß1.

(a)

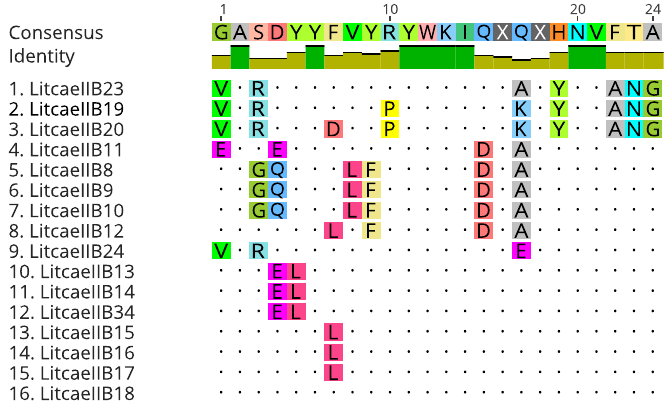

(b)

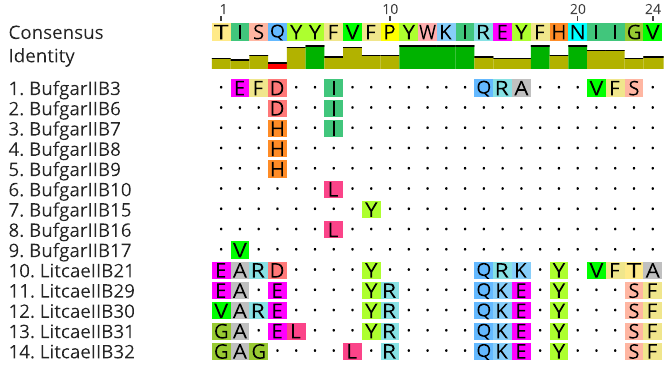

(c)

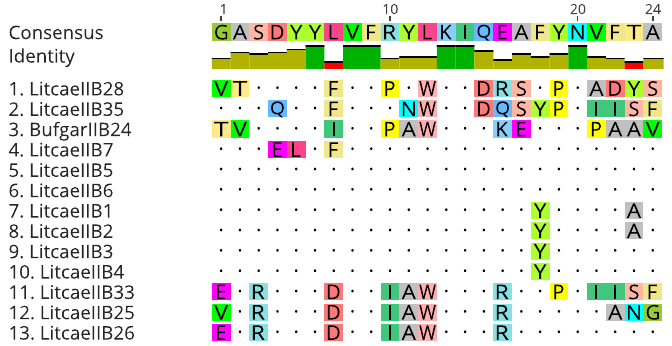

(d)

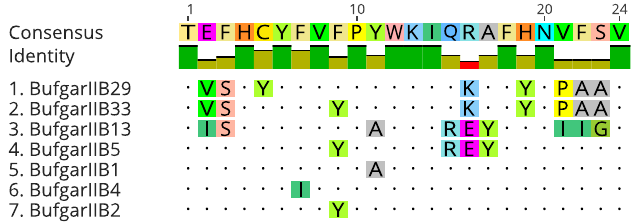

(e)

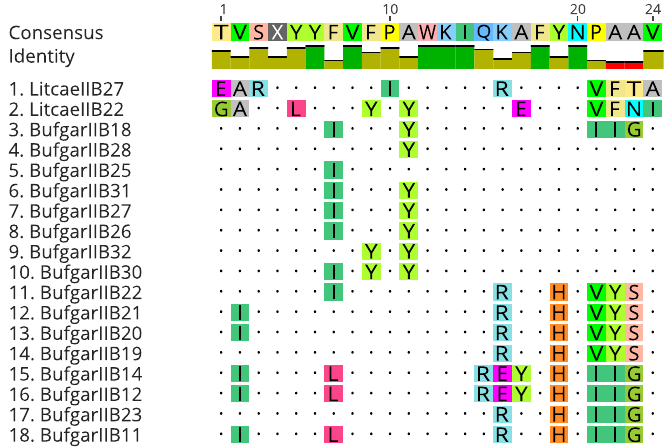

Figure S3 Alignments of amino acid for each MHC II supertype in Bd-resistant (*B. gargarizans*) and susceptible species (*L. caerulea*). (a) ST1, (b), ST2, (c), ST3, (d), ST4, (e), ST5.

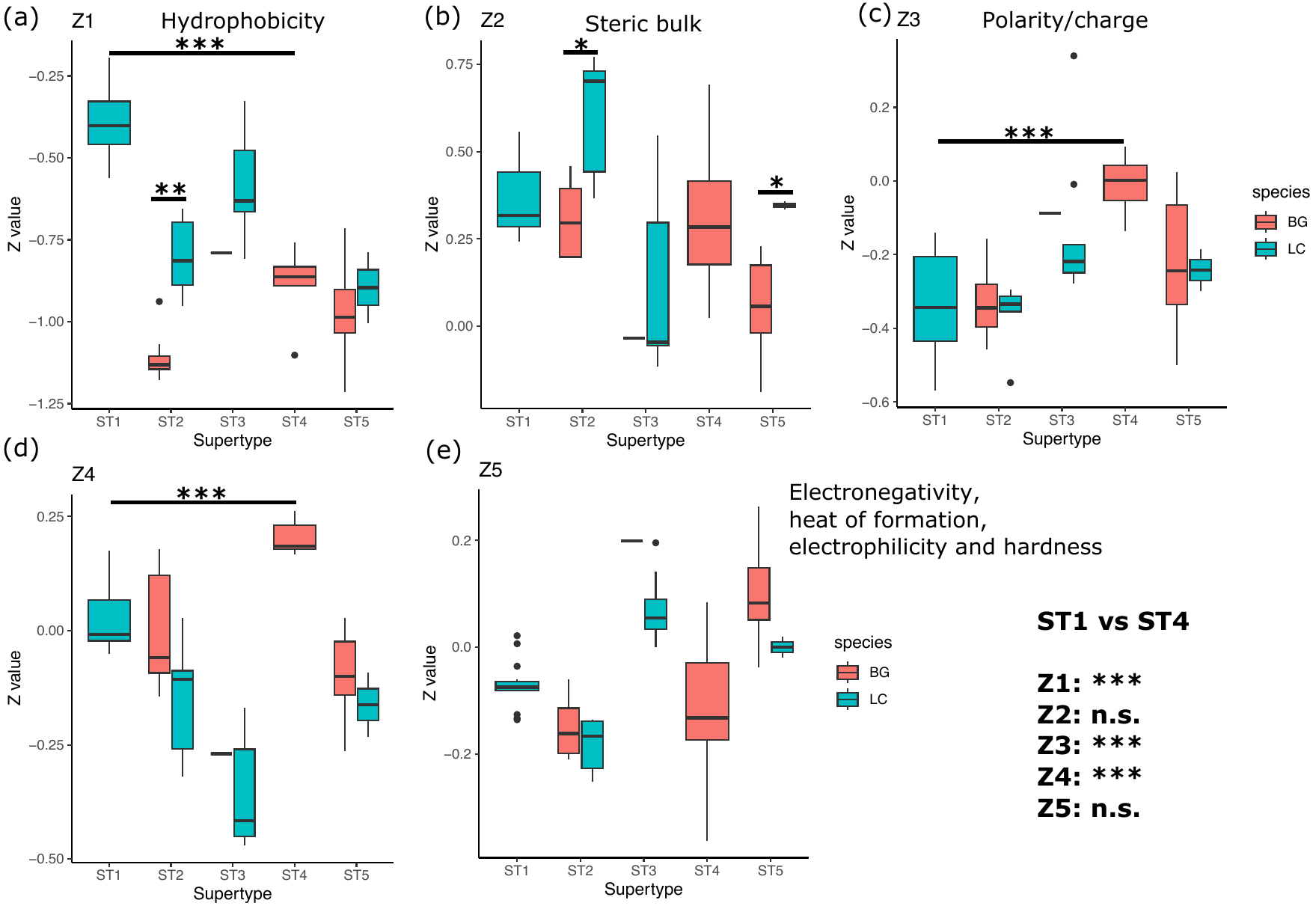

**Figure S4 Physicochemical properties of MHC supertypes in *B. gargarizans* and *L. caerulea***. (a), comparison of Z1 relating to hydrophobicity. (b), comparison of Z2 relating to steric bulk. (c), comparison of Z3 relating to polarity/charge. Comparison of Z5 (d) and Z5 (e), which related to electronegativity, heat of formation, and electrophilicity and hardness. When comparing between ST1 (unique supertype in *L. caerulea*) and ST4 (unique supertype in *B. gargarizans*), significant differences in Z1, Z3 and Z4 were detected (*p* < 0.001). ‘*’, *p* < 0.05. ‘**’, *p* < 0.01. ‘***’, *p* < 0.001. BG, *B. gargarizans*. LC, *L. caerulea*.

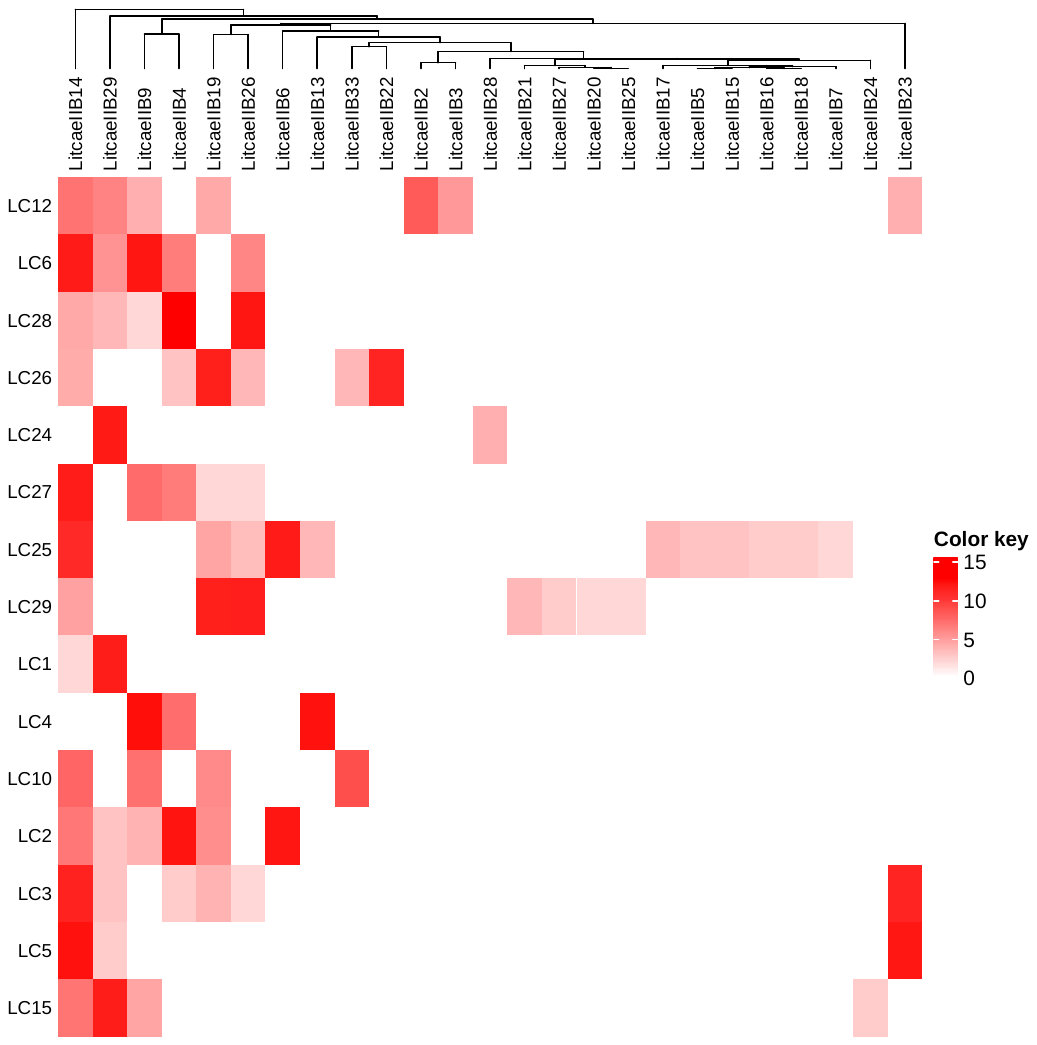

Figure S5 Gene expression pattern in Bd-susceptible and resistant L. caerulea. LC15 is the only Bd-resistant individual. The LC15 survived from Bd infection, is Bd-resistant *L. caerulea* and highly expressed LitcaeIIB29. However, individuals LC24 and LC1, highly expressing LitcaeIIB29, were susceptible to Bd infection. The resistance increases from top to bottom, with increasing surviving days after Bd infection.

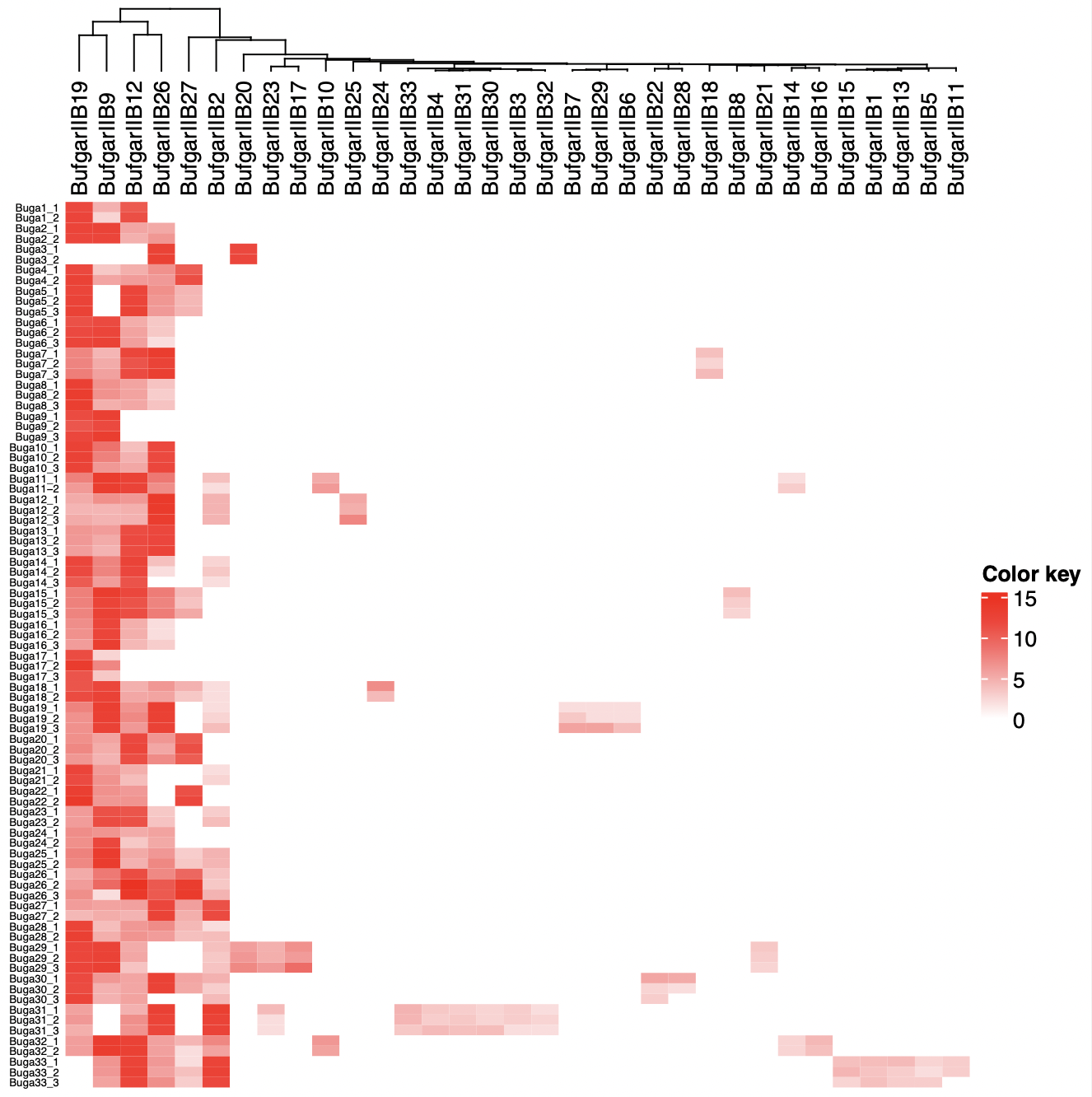

Figure S6 Gene expression patterns of MHC IIβ1 alleles across replicates in B. gargarizans**.** Distribution of 33 MHC IIß1 alleles and their relative gene expression across replicates in 33 individuals of *B. gargarizans*. Each color refers to a different allele. All the replicates for each individual were plotted in the heatmap.

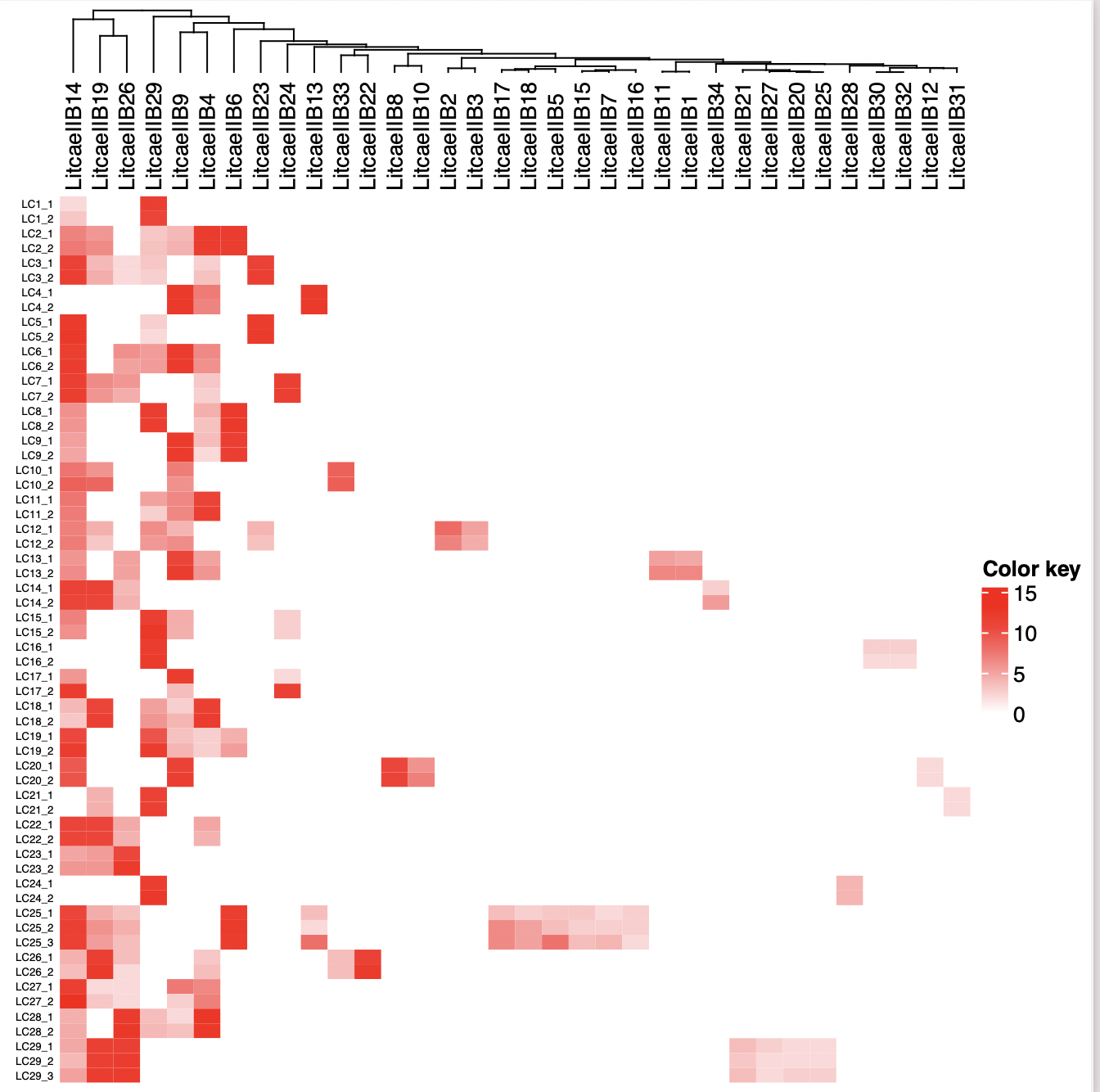

Figure S7 Gene expression pattern of MHC IIß1 alleles across replicates in *L. caerulea*. Distribution of 33 MHC IIß1 alleles and their relative gene expressions in 29 individuals of *L. caerulea*. All the replicates for each individual were plotted in the heatmap.

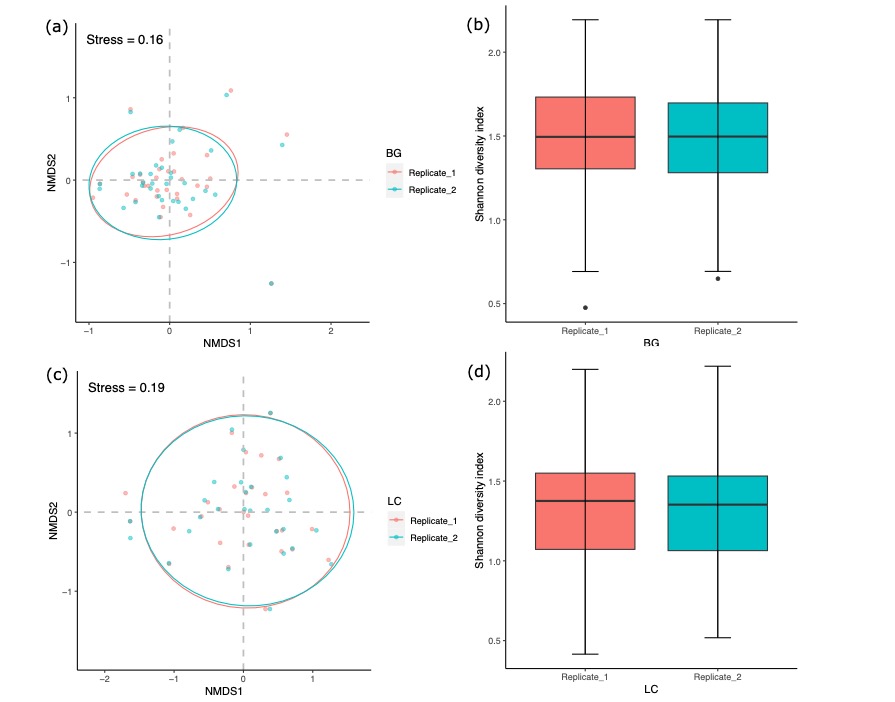

Figure S8 Comparing structure and Shannon diversity index for relative gene expression of MHC II alleles between duplicates in *B. gargarizans* and *L. caerulea*. Similar structure of MHC II relative gene expression between duplicates in *B.* *gargarizans* (a) and *L. caerulea* (c). Similar Shannon diversity of MHC II relative gene expression between duplicates in *B.* *gargarizans* (b) and *L. caerulea* (d). BG, *Bufo gargarizans*. LC, *Litoria caerulea*.

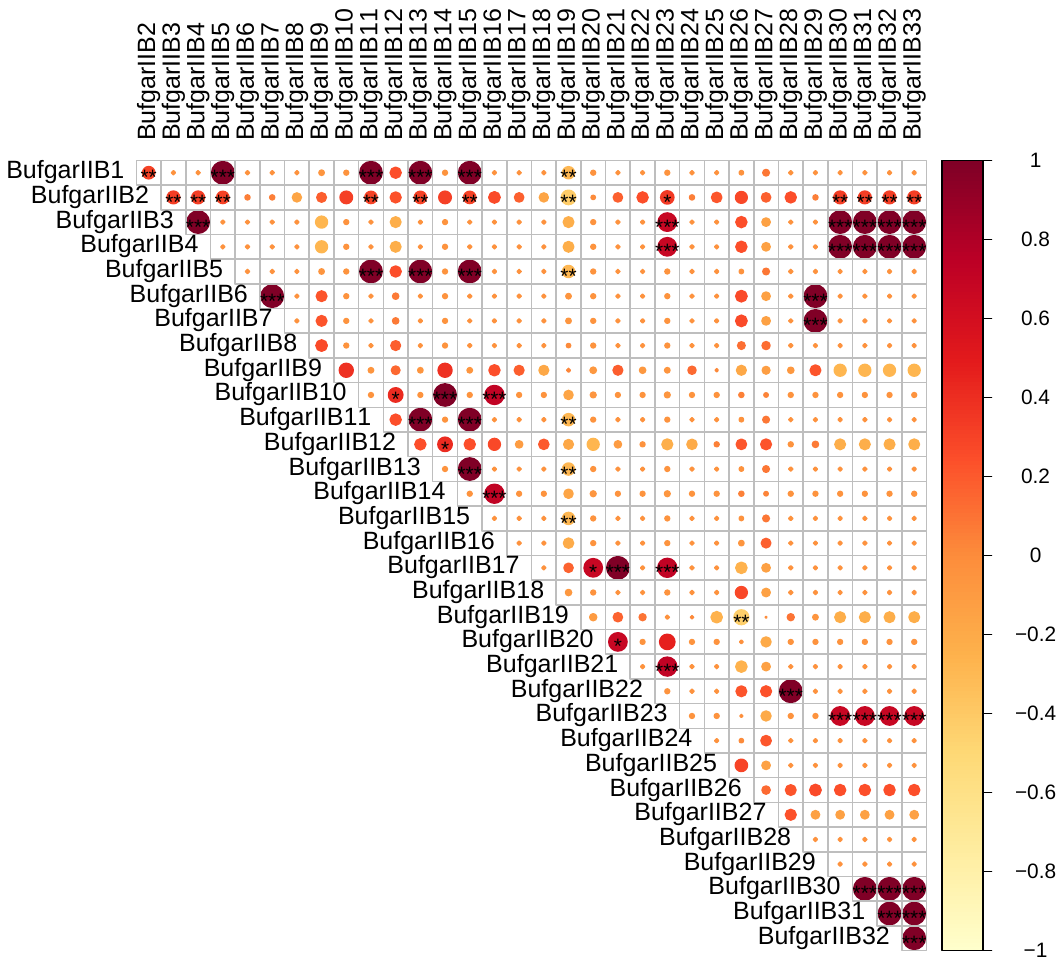

Figure S9 Pearson correlation matrix for MHC II alleles in B. gargarizans. Detailed correlation coefficients among MHC II alleles in B. gargarizans, showing significant positive and negative associations. ‘*’, *p* < 0.05. ‘**’, *p* < 0.01. ‘***’, *p* < 0.001.

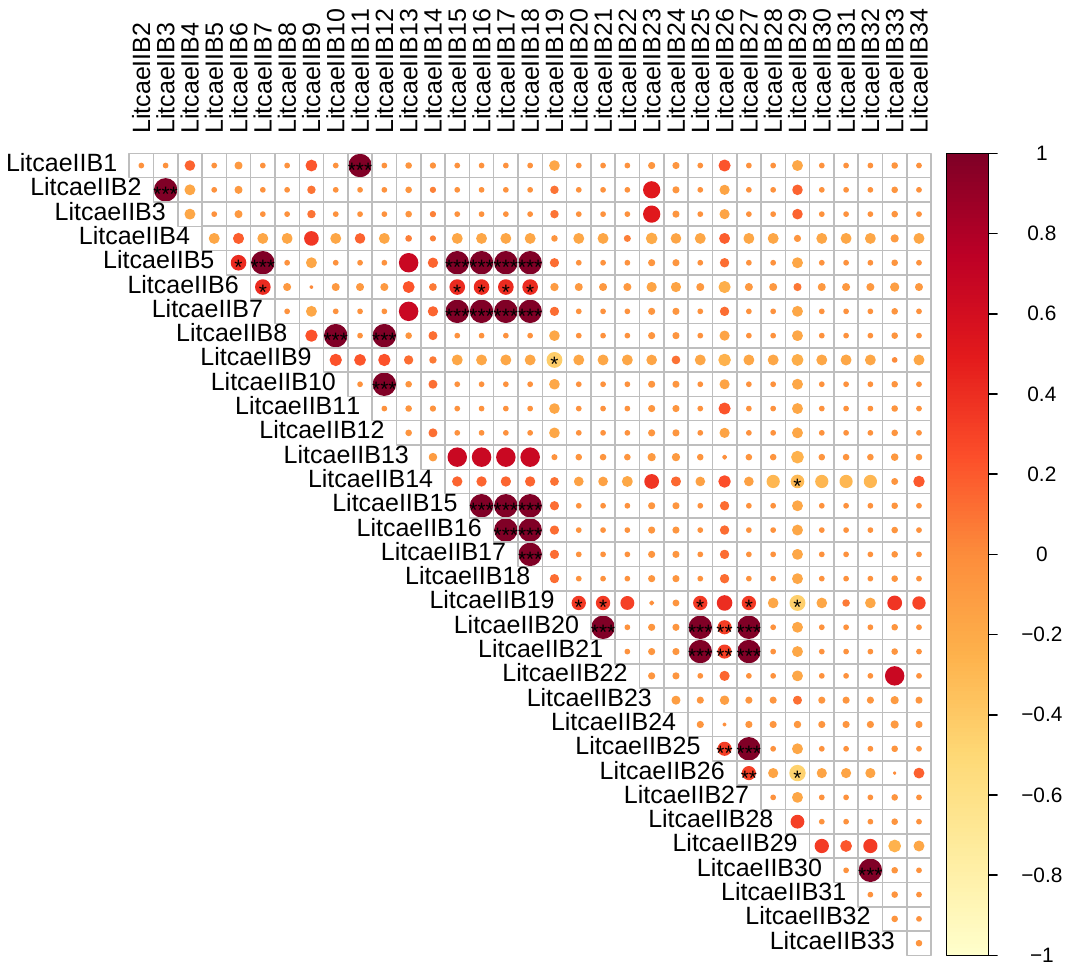

Figure S10 Pearson correlation matrix for MHC II alleles in L. caerulea. Detailed correlation coefficients among MHC II alleles in L. caerulea, showing significant positive and negative associations. ‘*’, *p* < 0.05. ‘**’, *p* < 0.01. ‘***’, *p* < 0.001.
